# Interactive Biology Cloud Lab Enables Authentic Inquiry-Based Science Learning at MOOC Scale

**DOI:** 10.1101/136317

**Authors:** Zahid Hossain, Engin Bumbacher, Alison Brauneis, Monica Diaz, Andy Saltarelli, Paulo Blikstein, Ingmar Riedel-Kruse

## Abstract

The Next Generation Science Standards (NGSS) and other national frameworks are calling for much more sophisticated approaches to STEM education, centered around the integration of complex experimentation (including real labs, not just simulations), data collection and analysis, modeling, and data-driven argumentation, i.e., students can behave like real scientists. How to implement such complex approaches in scalable ways is an unsolved challenge - both for presential and distance education. Here we report on the iterative design and large-scale deployment of an open online course with a “biology cloud experimentation lab” (using living cells) that engaged remote learners (> 300 students) in the scientific practices of experimentation, modeling and data analysis to investigate the phototaxis of a microorganism. We demonstrate (1) the robustness and scalability of the cloud lab technology (> 2, 300 experiments run), (2) the design principles and synergistic integration of multiple UI and learning activities and suitable data formats to facilitate NGSS-aligned science activities, and (3) design features that leverages the natural variability of real biology experiments to instigate authentic inquiry. This platform and course content are now suited for large-scale adaptation in formal K-16 education; and we provide recommendations for inquiry-based science learning in general.

## 1 Introduction

Inquiry-based learning is defined as “*an educational strategy in which students follow methods and practices similar to those of professional scientists in order to construct knowledge*” [31]. In authentic inquiry activities, students design and carry out experiments of varying complexity, formulate and test models, analyze and interpret their own, rich data and results. In the last few years, several national science learning frameworks, e.g., NGSS [8, 32, 44, 31], have been released, advocating to improve practices for STEM teaching and to make inquiry-learning more authentic. To realize this vision, a seamless integration of experimentation, data collection, analysis, modeling, and data-driven argumentation is needed. Without this integration, authentic science inquiry remains challenging to achieve [9], both for presential and distance education, especially when authentic real (instead of simulated only) experimentation is to be integrated [42]. Technologies provide promising means to that end, however, the design of technological supports for such complex integrative approaches is still in its infancy, especially when we consider the need for scalable, low-cost infrastructures. Various real and virtual labs in both presential and remote form have been developed and tested [18, 46]; each type having its distinct, context dependent advantages, and research is pushing to utilize their synergy [11]. At the same time, with online and blended education on the rise [45, 4, 25, 17, 41], it remains unclear how to integrate in internet-enabled learning systems experimentation-, modeling- and data analysis-practices based on real experiments and data at scale. Other existing remote labs have attempted to facilitate inquiry-based learning, but we are not aware of any that were designed to be accessed concurrently by many users at scale [18]. This paper addresses this issue and investigates how to enable and scaffold authentic scientific inquiry tasks remotely and at scale [24] while utilizing a new biology cloud lab paradigm [20].

This paper builds on a real-time and interactive biology cloud lab that we recently developed and successfully deployed in small, teacher-led classrooms [20]. Students in middle schools and in higher education ran interactive experiments to investigate how Euglena gracilis, a single-celled phototactic organism, senses and reacts to environmental stimuli such as light (Fig. 1). We demonstrated the potential of this cloud lab for NGSS-aligned science learning in classrooms [32] in ways that were previously not possible in biology education, not even on-site (given the limitations of passive microscopy, which is prevalent in K-12 education). One user study in [20] integrated experimentation with data analysis by letting students analyze the automatically-generated Euglena tracks quantitatively with Matlab. Another user study in [20] integrated experimentation with modeling by means of a modeling interface that enabled students to alter biophysical parameters in virtual Euglena and compare that data to real Euglena akin to bifocal modeling [3]. However, in our prior work, we have neither integrated experimentation-, modeling- and data analysis-based practices in a holistic manner nor deployed the system in settings larger than normal-sized classrooms and without an instructor being physically present.

**Figure 1:**
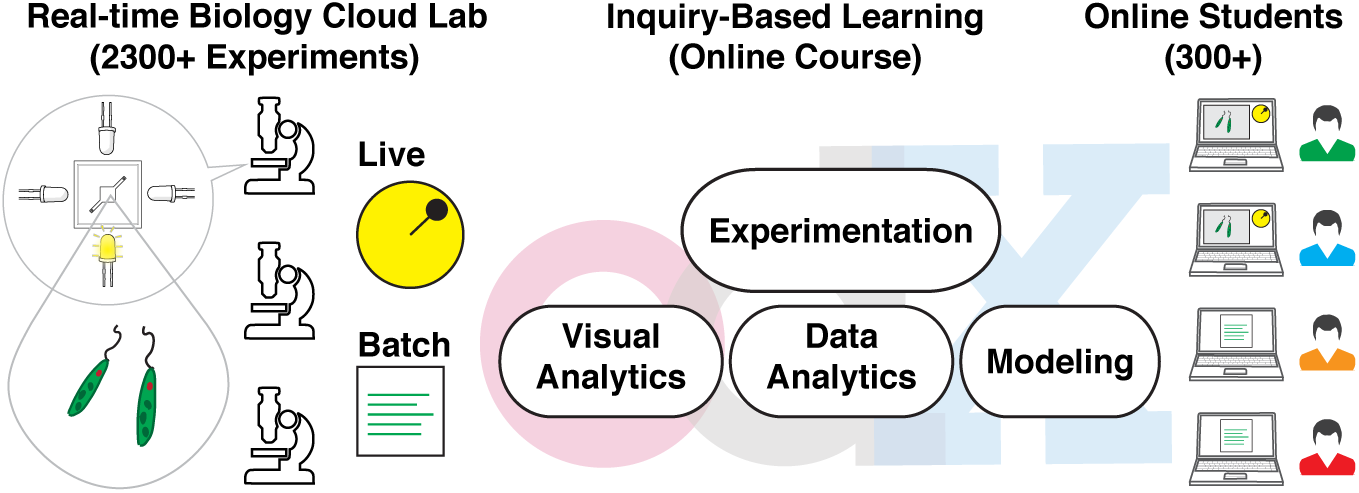
Integration of a scalable biology cloud lab into a MOOC (> 2, 300 experiments run by > 300 students). We explored the affordances and design rules of online experimentation science labs to enable inquiry-based learning.

In order to frame the research described in this paper, we start by reviewing some of the technological challenges for providing real experimentation at scale. Remote labs face more significant technological challenges of implementation than simulations, even more so when it comes to the life-sciences where noisy biological specimen need to be maintained in a functional state [21]. Consequently, a majority of existing remote labs have been developed for physics and engineering content [15, 18]. In addition, many are not robust or scalable enough for larger enrollments courses (MOOC-scale) as they are neither designed nor intended to be accessed concurrently by thousands of students. In our review, we found only one project that systematically tried to scale up an electronics lab in a MOOC environment [12]. This scaling problem motivates the distinction between *remote labs* and *cloud labs*. Here the former is more akin to remote computer sharing of a single instrument by one user [18], while the latter follows the modern cloud computing paradigm [16, 21, 20] that provides ubiquitous, on-demand access to a shared pool of configurable and distributed computing resources. The architecture of any given lab is likely to fall on some sliding scale between simple remote access and full realization of the cloud paradigm. Our biology lab followed this cloud paradigm: It is fully automated, low-cost, and scalable by design; the platform architecture load-balances concurrent experimental tasks with a cluster of back-end instruments (biotic processing units, BPUs) in a distributed and fault-tolerant manner, such that each BPU can run *∼* 100, 000 experiments per year for < $0.01 per experiment.

The main contribution of the present paper is to demonstrate that this cloud lab technology can support authentic science inquiry-based learning at large scale, and to distill design principles from the core technology, the user interface and the course for successful deployments of online labs and courses for inquiry-based learning. We used a popular online learning platform, Open edX [40], that integrates our cloud technology to deploy a short course (on the scale of 4 hours) that engages students in scientific inquiry. This required the use and further development of existing as well as novel novel user interfaces and technologies so that students could design and execute experiments of varying complexity, model scientific phenomena, analyse, and interpret the obtained data. We report results from the iterative design process and public release with over 300 users. We primarily concern ourselves with the course design and the related HCI technologies that enabled the fundamental activities for inquiry-based learning to occur through the Internet at scale, i.e., online students are enabled to perform activities similarly as real scientists would do. A thorough investigation regarding the learning outcome due to this cloud lab and online course is planned for the future.

We organized this paper in the following logical sections: First, we review inquiry-based learning and discuss all its key phases. We introduce the biological phenomena, phototaxis of Euglena gracilis, which is the central learning theme of the course. We then discuss the key technological improvements compared to our previous cloud lab [20] and how we integrated multiple HCI components to allow a larger scale deployment in a MOOC environment; we then also estimate the total capacity and throughput of this improved cloud lab implementation. For each of the seven units of the MOOC, we discuss content layout, scaffolding and the HCI design principles that facilitates inquiry-based learning; these design principles were derived by analysing student logged activities and feedback during the multiple course iterations and from previous pilot studies. We present a case study to demonstrate how a single user experienced inquiry-based learning as she went through the course from the start to the end. We assess the outcome of this MOOC deployment using students’ logged activities and voluntary feedback along multiple dimensions. Finally, we summarize lessons learned and discuss future work including a potential path towards massive deployment in formal school contexts.

## 2 Rational for Research Design, Online Course Layout, and Technology Implementations

We adopted an iterative design-based research approach [1, 14] to develop, deploy, and evaluate an open online course for inquiry-based science learning. We chose the format of a mini-course with the intended student effort of *∼* 4h over one week. This allowed us to rapidly iterate course features based on study outcomes over multiple week-long course offerings. We refined our cloud lab technology and user interfaces by incorporating lessons from our previous pilot studies [20] in order to accomodate users at MOOC scale. We implemented this course in the Open edX framework [40] with a diverse target audience that ranged from middle-school to university students and science teachers. To engage students in all the key phases of the inquiry-based learning (Fig. 2A) in the context of learning about the target biological concept - *phototatic* behavior of Euglena cells (Fig. 2B) - we were required to holistically combine all the experimentation, modeling and data analytics UIs (Fig. 3) within a single cloud lab platform. We will now describe in detail these three key facets of the current work: 1) inquiry-based learning, 2) the target biological concept (*phototaxis*), and 3) the underlying cloud lab technology with its interface and interaction design.

**Figure 2:**
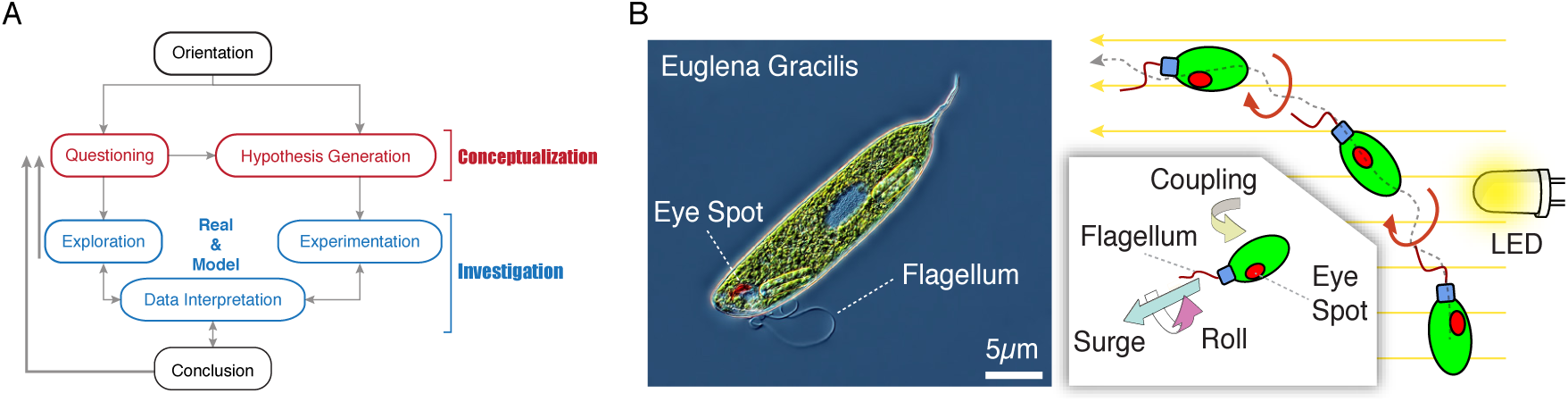
Scientific inquiry and target biological concepts of the online course: (A) The course takes students through the different phases and sub-phases of a full inquiry cycle in an integrated manner. We adapted the schematic from Pedaste et al. [31] with emphasis on exploration and experimentation both with real specimen and models. (B) Euglena Gracilis is a single-celled organism that performs negative phototaxis, orienting and moving away from light, by rolling around its long axis but also turning sideways via a feedback coupling between the eye spot and the flagellum.

**Figure 3:**
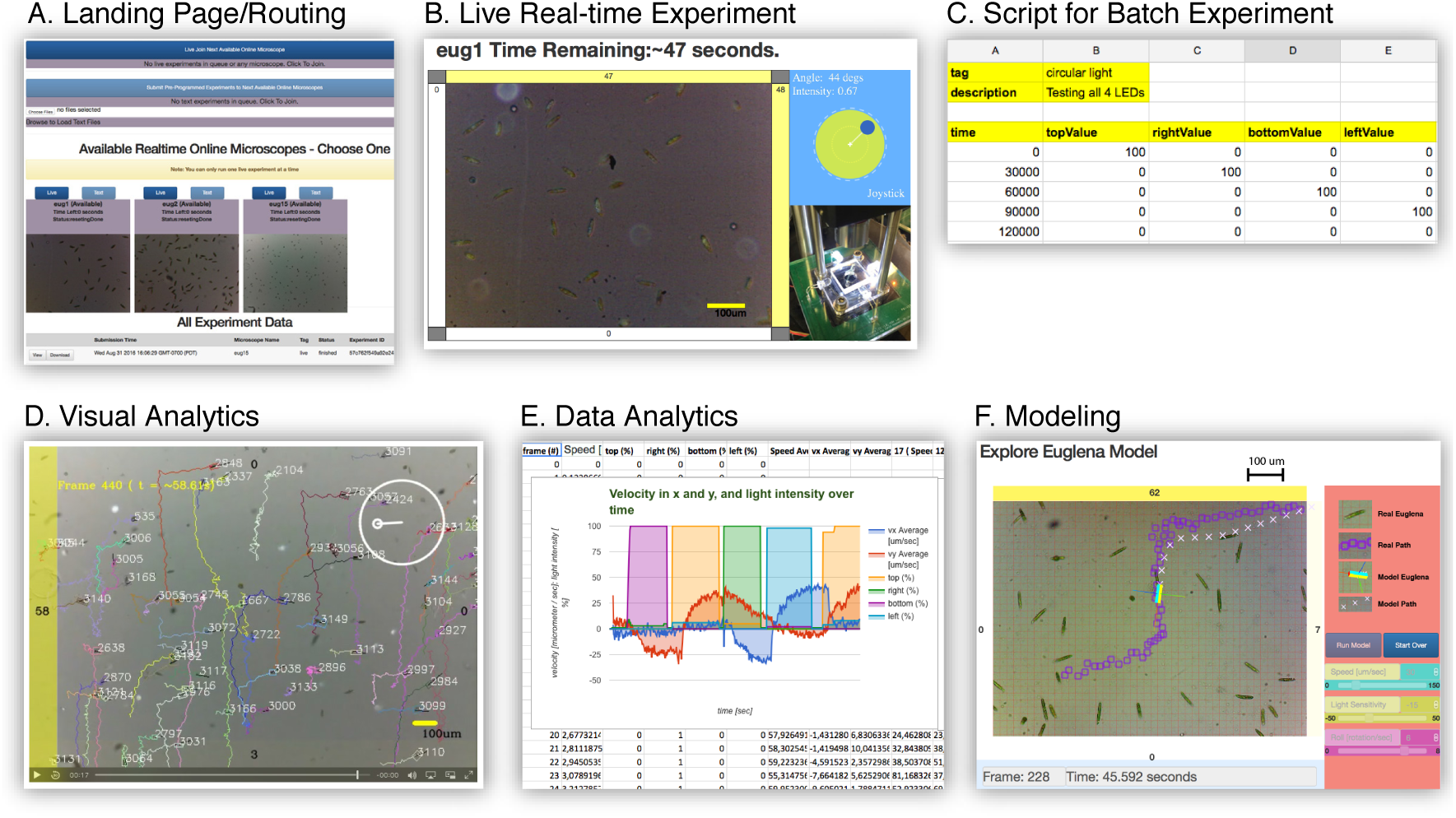
HCI modules to deliver procedural learning goals of the course in an unsupervised MOOC setting. (A) Landing page to route students among a suite of online microscopes. Students can either choose to get auto-routed to the best microscope or choose a specific one. (B) Realtime Euglena biology lab in *live* interactive mode. (C) An experiment script in CSV format for the *batch* mode. (D) A playback movie viewer for visual analytics via automatically tracked Euglena cells. (E) Google Sheets application for data analytics, including statistical analysis and graphing of Euglena traces. (F) Modeling applet simulating Euglena overlaid on pre-recorded video.

First, we turn to the phases of inquiry-based learning as defined by the NRC [39], which also defined learning goals and best practices for science labs: (1) posing questions and formulating testable hypotheses, (2) designing and carrying out investigations, (3) using tools to make observations, gather and analyze data, (4) building, evaluating, testing or verifying explanatory models in light of empirical data, (5) interpreting and communicating results. In order to coordinate and contextualize these phases, we based our course design on a recently synthesized model of the inquiry cycle [31] (Fig.2A; Table 1 column 2). However, without proper scaffolds and guidance, inquiry-based activities are hardly effective for learning and can overwhelm students [24, 46]. Units #1 to #5 are therefore structured to guide students through the inquiry cycle with a given set of questions and investigations to perform: students started with passive observations to conceptualize the problem and moved on to active experimentation, engaged in qualitative data analysis, proceeded to model exploration and parameter fitting, and then advanced to quantitative data processing and graphing. In unit #6 this then culminated in a self-guided project where students formulate and investigate their own hypothesis. With each phase, this course structure introduces a new corresponding user interface (UI) tool (Fig. 3), while providing multiple opportunities to revisit earlier activities.

**Table 1:** Final course layout. Note that each unit essentially introduces a tool (“instrument”), and engages students in a different inquiry practice with a new bit of biological content. ^*†*^In unit #6, students were asked to postulated testable hypotheses about Euglena phototaxis, but to pursue the actual experimentation and analysis was left optional.

| Unit # | Scientific/Inquiry Task | Target Biology Concept | Technology |
| --- | --- | --- | --- |
| 1 | Passive observation | *Euglena* are single cells | Online microscope |
| 2 | Active experimentation | *Euglena* respond to light | Real-time interactive & batch experiments |
| 3 | Visual analytics | *Euglena* roll around their axis while swimming | Post-experiment video analysis |
| 4 | Exploration and conceptualization of models; parameter fitting | From structure to function: feedback loop between eye spot and flagellum | Modeling Applet |
| 5 | Data processing; interpreting graphs | *Euglena* speed does not change (much) when light is turned on | Google Sheets |
| †6 | Self-guided project; generate and test hypotheses | Learn more about *Euglena* phototaxis | All the previous tools as necessary |
| 7 | Summary, reflection and feedback |  |  |

Second, the target biological concept of the course revolve around Euglena phototaxis [13] (Fig. 2B; Table 1 column 2). It exemplifies the general taxis principle applying to many cell types and stimuli, e.g., Euglena gravitaxis [26] or bacterial chemotaxis [2]. Euglena has an eye spot that senses light coming from one direction only; furthermore a flagellum that allows the cell to swim forward and rotate around their axes. The photoreceptor is coupled with the beating pattern of the flagellum. Appropriate coupling of strength and directionality allows the cell to stably swim toward or away from the light. As to be expected from biological systems, not all cells behave exactly the same, i.e., these microscopic cells exhibit variability and individuality. This biological noise and variability merits particular consideration for this cloud lab, as it can interfere with a consistent user experience. At the same time, dealing with real experimental data with natural variability can also be a very productive learning experience [3], and was identified as a key laboratory experience by NRC [39, 43]. In this course, we used a sequence of target biological concepts starting with basic cell behaviors and progressed to more advanced ones like feedback regulation and noise, and ultimately encouraged students to embark on a self-driven research investigation as an optional final project.

Third, for an effective engagement with the inquiry activities, students must be at ease with various instruments and user interface tools to execute experiments (free form exploration as well as controlled), explore models, collect experimental data, and infer results via visual and data analytics (Fig. 3; Table 1 column 3). We had to centralize all these tools in a way that reduces extraneous effort and switchingcost between phases and sub-phases of the inquiry cycle (Fig. 2A), thus providing a seamless *laboratory experience* [39]. Compared to our previous work [20], we made significant additional technical and HCI strides. First, we increased the experimentation throughput of the system by over two-fold by making all the backend server system asynchronous with respect to each other. This was crucial for a large scale deployment such a MOOCs as it cut down the average wait time of students by half. Secondly, we made crucial HCI advancements to make the system more accessible to a broader audience as discussed in the following. We adopted ubiquitous file formats such as CSV and MS Excel for programming experiments and experimental data exports respectively (Figs. 3C,E). The MS Excel format can also be imported into the freely available software such as Google Sheets, which opened up our system to a much broader audience compared to the Matlab-based data analytics interfaces in our previous pilot studies [20]. The movies resulting from the experiments were augmented with various visualization elements (Fig. 3D). We adapted our previous modeling and parameter fitting interface [20] using a predetermined stimulus sequence instead of joystick-induced stimuli to reduce cognitive load (Fig. 3E). For the more technical details of the cloud lab system, we refer readers to our previous work [20].

We implemented the course with 7 units (Table 1). Each unit introduces students to a new *inquiry phase* (task) (Fig 2A), a new *biological target concept* (Fig. 2B), and a new HCI module (Fig. 3). Units were designed to take 20-60 min each. The unit #6 encourages students to postulate a testable hypothesis and voluntarily pursue a self-guided research by going through the whole inquiry cycle on their own. Unit #7 provides a summary and collects students’ feedback. We purposely made exposure to biological variability a central theme of the course content. Before launching the course, we iteratively tested and updated this course with 1 – 3 students at a time for a total of *∼* 20 students, with the goal to optimize the progressive increment of the complexity for the learner and also make the course duration as short as possible, leading eventually to the week-long course layout described in Table 1.

## 3 Course Deployment at Scale, Iterative Refinement, User Study Results, and Design Lessons

The course was repeatedly offered six times (six weeks) with minor updates between successive offerings (Fig. 4A). Students were recruited via monthly Open edX newsletters; additionally, 300 teachers were contacted directly. (We refer to all participants as students from now on.) A total of 993 students signed up (sessions distribution: 97, 259, 296, 157, 76, 48), of which 325 (35%) started the course and created an account in our cloud lab. The completion rate was 33%, (108 students) which we based off of the students who had answered at least one question in every unit (except for unit #6, which was partly optional). Note that this is a much more conservative estimate compared to our work-in-progress paper [19], where we only counted the number of students answering a question in unit #7 compared to unit #1. Students came from 46 countries (Fig. 4B), mostly from the USA (42%) with a median age of 32 (IRQ=19), 47% of whom were female. Students took 3.5 ± 1.1h (Mean ± Stdev: we will use this notation throughout the paper unless stated otherwise) to finish the course, with each of the seven units taking ∼ 30 mins to complete, except unit 5, which took ∼ 1h.

**Figure 4:**
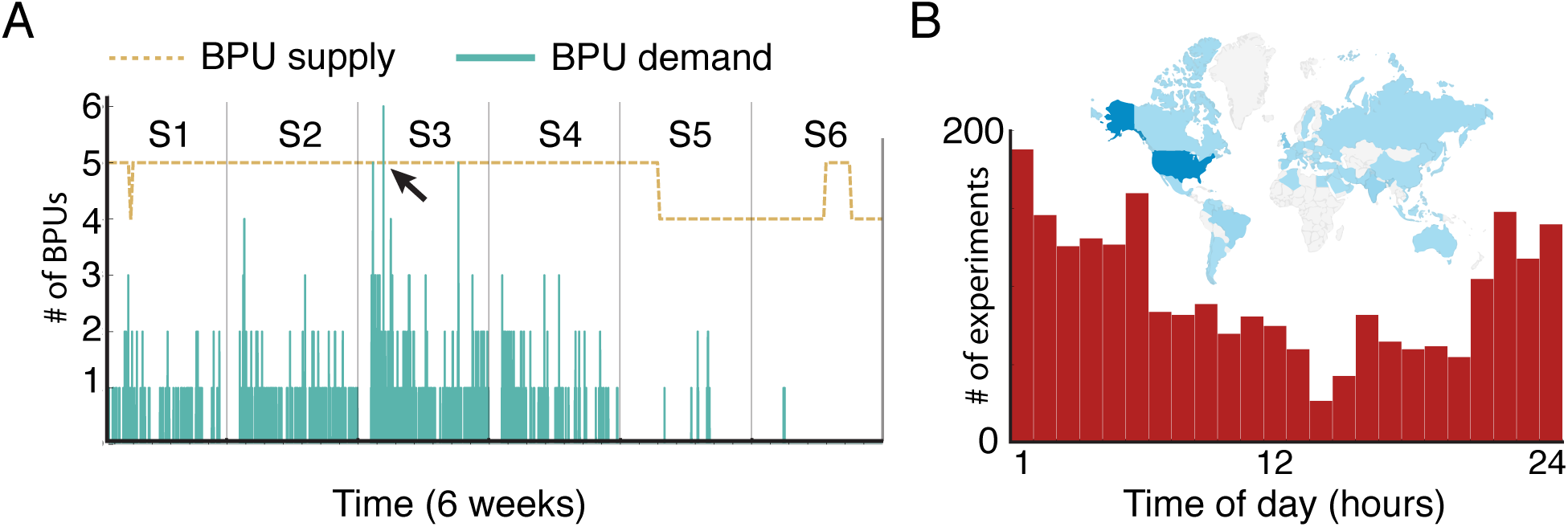
Long term usage of the cloud lab: (A) BPU (online microscope) demand during the first 6 weeks (S1,S2 and so forth) was well below the supply at all time except for a singular incident (marked with arrow) when a student had to wait for her turn. (B) System access pattern over 24h period during 6 weeks. The traffic was mostly amortized over the entire day with peak activity seen during the middle of the night (PST). The inset shows the origin of the traffic based on countries with the US providing most students.

Students were able to run experiments in both interactive *live* mode (Fig. 3B) (1 minute long) and preprogrammed *batch* mode (Fig. 3C) (capped to maximum 5 minutes long). In live *live* mode, students would interact with Euglena in realtime using a software joystick with a live video stream feedback, while in the *batch* mode students would submit a preprogrammed script (in JSON or CSV) of a light sequence, instead of using the joystick, to be executed offline without any live video feedback. However, in all cases, a timelapse video of the experimental session is recorded and made downloadable along with other data. A total of over 2, 300 experiments were executed, with 7 experiments on average per student (students who completed the course ran 12 ± 6 experiments versus students who did not complete the course ran 5 ± 6). During these 6 weeks, students hardly experienced any wait-time (median: 4.8 secs, IRQ: 1.55 secs, which is within the system loss time for routing) for *live* experimentation as the demand for the online microscopes was always below the supply, except for a single incident (Fig. 4). Due to the worldwide accessibility, the usage of the cloud lab was effectively amortized throughout a day, 7 days per week (Fig. 4) though we observed peak traffic typically in the middle of the night (based on the time zone were most students were situated, i.e., US).

The cloud lab scales linearly with the number of BPUs, and the demand characteristics depend on the deployment scenario, e.g., a week-long MOOC versus an hour-long classroom session. In this work we only concerned ourselves with a MOOC deployment with worldwide access, and assumed that the cloud lab was used continuously throughout. During the 6 weeks of the course, we had 4 − 5 operational BPUs (Fig. 4A). The total available capacity of the system was 1, 800 experiments per week with 4 BPUs (at a rate of 2 experiments per minute and discounting the 8 self-monitoring experiments per hour. See [20] for details but note that the throughput of our current work was improved by 2×. We saw maximum traffic on the third offering (week 3) of the course with 1, 000 experiment being executed by the students. Therefore in terms of raw capacity, our system could have conservatively supported 10× more students assuming an effective system utilization of half a day. This translates to a conservative estimate of > 500 students/week across the globe (10× of 54 active students/week on average -325 in 6 weeks). Even though the current work does not concern itself with a school deployment, we estimate that 12 BPUs are sufficient for this purpose: Euglena inside the BPUs are in a responsive state 61% of the time (empirically measured [20]), e.g., 7 BPUs are readily available for a typical class of 36 students working in pairs. Experiments take 1 minute, hence each student pair could start a new experiment every three minutes, with time in between for analysis and planning. Over a whole class period of 45 minutes each pair then could run more than 10 experiments, which is sufficient for a typical content progression (see analysis further below). Lastly, the manual effort to maintain the cloud lab on the backend was < 1 h/week to exchange organisms and microfluidic chips.

We will now discuss the design of each course unit (see Table 1), and the design rationale of the corresponding HCI module and technology, analyze users’ activities and feedback, and derive Human Computer Interaction (HCI) design principles. The course contents design and all the HCI design principles were derived through an iterative design process via several pilot studies, including our previous work [20], user interviews and pilot studies. The generalized HCI design principles in each unit are applicable to other cloud labs with real experimentation, even beyond biology, with the primary goal to support students’ successful engagement in inquiry practices. Unit 7 focuses on user feedback and therefore does not relate to any specific design aspects. All the statistics in the following sections are based on the logged user interaction data, pre/post interview questions, voluntary student feedback and performance on certain course exercises. We automatically logged details of every experiment, such as time of submission, time of execution, joystick movements (*live*), programmed instructions (*batch*), while the timelapse videos produced by every experiment were available on our server for download.

### Unit 1: Observation (online microscope)

Students were introduced to the cloud lab dashboard interface (Fig. 3A), i.e., how to select an online microscope (BPU), how to observe the Euglena cells in real time, and how to watch the resulting experimental video afterwards either directly on the website via streaming or after downloading. Students were tasked to describe their observations in a free form manner. We deliberately started with a passive observation, rather than asking students to already explore the light responses, as earlier pilot studies [20] and interviews with users as part of the iterative design process have shown that premature interactivity without proper foundation being established could overwhelm students, especially when working with a noisy biological system. This passive observation was then followed by a short description on the basic biology of Euglena: they are photosynthetic organisms and they can detect light using an eye-spot; there was no mention of phototaxis yet. During the first course offering we found that users with low Internet bandwidth struggled with the *live* lab. In later offerings we therefore alerted students about the Internet bandwidth issues upfront, and emphasized about the option to download the stored experimental video to watch offline. We noticed that many students with even 5 Mbps of Internet bandwidth were able to successfully finish the course.

#### Design Rationale

The cloud lab dashboard UI provided two methods for selecting an online microscope: 1) auto-select a BPU based on the best available one in terms of Euglena responsiveness, which was automatically monitored by the backend system, and estimated wait-time [20], and 2) manually select a BPU, e.g., in order to do repeat experiments on the same device. We also implemented an external camera that provided students with a view on the microscopes in the cloud lab in order to emphasize that these were real microscopes (Fig. 3B). This was motivated by several previous studies [7, 6, 23, 28] that have demonstrated increased student motivation when they perceived that the lab activities were indeed real.

#### Student activities and feedback

Student found the online microscope with real organism to be very useful (7.5 ± 2 on the scale of 0-9, N=52), and voluntarily provided positive comments during the optional feedback session, e.g. *“I liked the focus on one organism… I liked the use of live microscopes to study Euglena.”* 100% of the students were able to correctly identify Euglena in a follow-up question where they had to choose which one of a set of pictures showing various microorganisms related to what they had observed in the online microscope. Students answered questions about the quality of live streaming and download speed: 40% percent of students reported *“really nice”* (likely due to the Internet speed); another 45% reported that it was *“at least reasonable.”*.

#### General Design Principles

Students should be enabled to first immerse into the real aspect of the cloud lab, which is achieved through realtime observation of the underlying scientific subject (Euglena in this case). The system should also provide an external camera view of the actual workings of the lab even if that camera does not have any direct consequence on the experimentation, but to increase the credibility of the system. Furthermore, the system should provide a timelapse recording of the experiments, along with other data, so that students can investigate offline at a later time, which also helps mitigate lower bandwidth issues.

### Unit 2: Experimentation (Interactive and Scripted)

Unit 2 introduced students to the interactive joystick to interact with the online microscope (Fig. 3B) to actuate directional light stimuli. Students were prompted to then run experiments to explore how Euglena reacted to light stimuli. We primed students with simple test questions, e.g., *“In which direction does the light shine when you pull the joystick in this direction?”* to eradicate misinterpretation of the instrument usage early on. In this unit, we also introduced the *batch* mode experimentation as an alternative approach to running experiments (Fig. 3C).

#### Design Rationale

The goal of the *live* interactive experimentation was to allow students to intuitively calibrate themselves to the light behavior, the time of reaction and the length scale of the Euglena biology in an easy exploratory manner. The Euglena phototaxis with respect to light is non-linear, has an implicit time delay (takes approx. 7 secs for the swarm to respond to light), and only visibly activates when the light intensity is above 40% [20]. In a previous iLab pilot study [20], which was only based on *batch* experimentation, we noticed that students had great difficulties in bootstrapping experiments with the right timing and lighting condition to induce negative phototaxis. Realtime interaction helps students establish an intuitive sense of the various scales, while *batch* experimentation might be better suited for further controlled investigation. The *batch* mode also enables students with low Internet bandwidth to participate with the offline timelapse viewer. Biological systems, unlike physical systems, often undergo unpredictable natural variability (e.g., cells dying, culture contaminated, or population undergoing circadian rhythm), which makes repeatable and consistent experimentation challenging. To mitigate that, we implemented auto-monitoring and self-correction of the biological state for each BPU [20], and then routed students automatically to the optimal BPUs. On the other hand, the moot point of a cloud lab with real biology from an educational stand point is to expose these very natural variability for a real life scientists experience, which we initiated by asking students to repeat their experiments on at least two different, manually chosen microscopes.

#### Activities and Feedback

Prompted by the open-ended question: *“What do you see?”*, 83% (N=163) of students reported Euglena responded to light, among which 62% recognized negative phototaxis, 10% positive phototaxis, 7% both types of phototaxis and 4% *“spinning”* without linear motion. These observations indicated that the Euglena light responses in the experiments were clear enough for the majority (negative phototaxis is the expected dominant response), and most students were able to self-*“discover”* negative phototaxis in light of biological variability. The remaining 17% of students did not give the expected answer, and we identified in the students’ experimental data multiple reasons: some students used too low light stimulus or did not wait long enough for Euglena to respond, whereas some students self-selected a microscope with either too few cells on the screen or cells that were in a state of too much light sensitivity, with higher likelihood for cells to just spin on the spot. In later course offerings we therefore asked student to run two additional experiments on different self-selected microscopes, which successfully mitigated these issues, furthermore emphasized that each instrument is different. Students mainly used *live* experiments (2255 *live* compared to 69 *batch*). Students rated this interactive experimentation to be very useful (8.4 ± 1.5 on the scale of 0-9, N=52) and expressed enjoyment, e.g. *“The ability to see real Euglena and interact with them (using the LED lights) was really interesting. The real-life interactions made this course much more fun. I thought it was neat to be able to actually control the parameters and experiment with the lights.”*

We also analyzed the types of light stimulus experiments run by students (Fig. 5). For every experiment, we collapsed the joystick positions over time into a single image by convolving a Gaussian distribution on every joystick position and performed hierarchical clustering on 1321 experiments (72% of all *live* experiments that had sufficient mouse movements). A judicial cut-off in the hierarchy revealed six dominant clusters (silhouette score [35]: 0.53 in 0 – 1 range; see inlets in Fig. 5). The corresponding cluster centers correspond to experiments that either focused on directional responses (C4, C5, C6) or on responses to varying light direction (C1) or intensity (C3). A Locally Linear Embedding (LLE) [36] analysis with two components (Fig. 5) reveals the general spread and subtle variations among the experiments. Note that in many experiments students tested intermediate light intensity values (Fig. 5 C2 and C3) as opposed to full intensity. This analysis demonstrates that students have freedom of performing different types of exploration and assess different variables (direction, intensity, duration), and that the joystick provides an intuitive input; this analysis also points to the potential for future analysis using learning analytics and data-mining techniques regarding student experimentation strategies.

**Figure 5:**
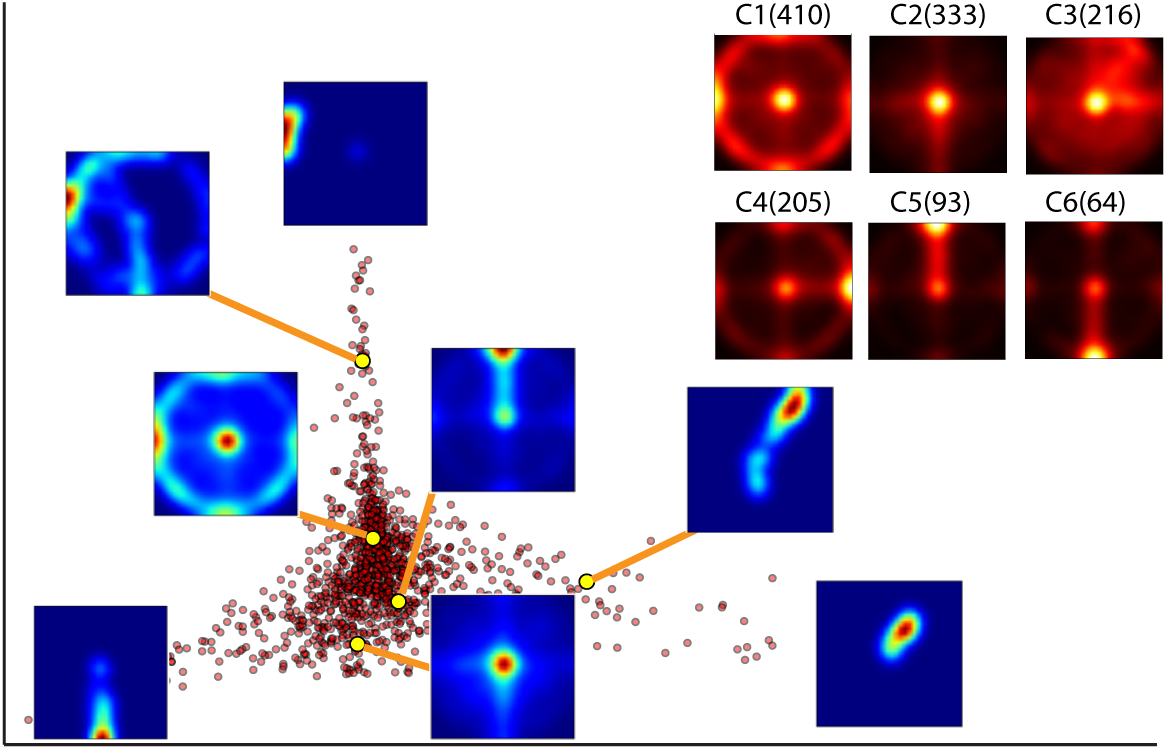
Typical *live* joystick experiment run by the students and a Locally Linear Embedding (LLE [36]) analysis shows the varieties among all the experiments run before unit #3 (N=1321). (Note: The axes are not labeled on purpose as they bear no semantic meaning in LLE, which is a non-linear dimensionality reduction technique to mainly show the spread in the dataset.) The inset shows the six dominant clusters extracted using hierarchical clustering (silhouette score of 0.53 in 0-1); numbers of experiments in each cluster in brackets.

#### General Design Principles

The course layout and UIs should allow students to both “play” with the biology in a free form manner using simple and intuitive interfaces in *live* mode, but also provide the possibility for programming experiments in *batch* mode that allows for better controlled experiments, and which also mitigates challenges with low Internet speed. A cloud lab platform, especially for biology, should strive to allocate experimentation equipments (BPUs) that have higher signal-to-noise ratio (e.g. through auto-monitoring BPUs), thus providing a clearly observable and repeatable experience. On the other hand, the platform should leverage the real cloud lab to expose the biological variability of the system to facilitate rich exploration based educational experience. Letting students run experiments on different (but seemingly equivalent) instruments further emphasizes this concept. It is also important to ensure that the students fully understand their inquiry instruments, rather than just focus on their object of study.

### Unit 3: Visual analytics/Qualitative data interpretation

Unit #3 instructed students to analyze and explain their movie data more closely (Fig. 3D), where Euglena exhibit a wobbling, meandering motion as apparent via the overlaid visualizations of their tracks (Fig. 6). Students then performed simple, direct measurements regarding speed and rolling frequency - solely based on visual analytics using the overlaid tracks, the timer and the scale bar.

**Figure 6:**
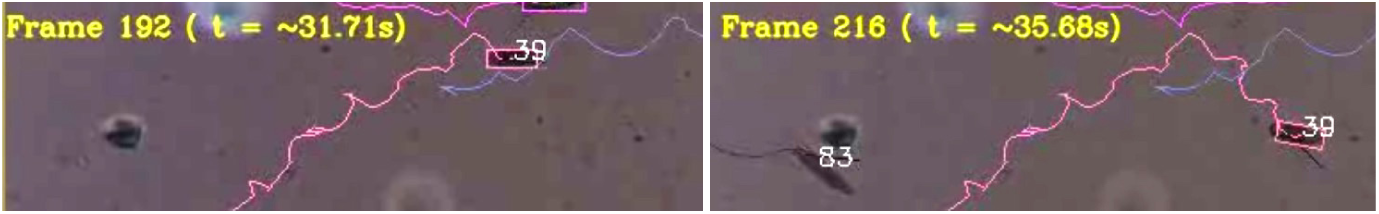
Post-experiment movie analysis and back-of-the-envelope measurements. Example shows the meandering swimming path of Euglena cell #39 in two different image frames; measuring the distance provides the forward speed of the cell, counting the peaks provides the rotational speed of the cell about its long axis.

#### Design Rationale

The visual analytics component was an essential new UI addition, as during our previous work [20] and during our pre-studies some students did not recognize negative phototaxis or the wobbling Euglena motion easily. We therefore generated movies that contained all possible information overlaid on the original movie (Fig. 3D), i.e., a scale bar, timer, frame number, side bars indicating light direction and intensity (both by intensity of the bar as well as in numbers, i.e., aiming for redundancy whenever possible), a joystick animation, and finally the cell tracks, where each cell had a unique ID. This ID could be cross-referenced with a downloadable data file (discussed in unit# 5). We rationalized that students would be enabled to recognize features that might otherwise go unnoticed, such as the side-way wobbling of the cells. Furthermore, students could measure several quantities by simple inspection, e.g., size of Euglena, speed, response lag time due to changes in light by inspecting when the tracks start to bend, or rotational frequency of Euglena rolling around its long axis by counting peaks of the path due to the wobbling motion. These activities were also intended to prime the students for the modeling exercises in the next unit.

#### Feedback

50% of the students hypothesized that the wobbling would allow for better detection of light direction (which we considered to be correct); other answers included better maneuverability or faster escape from predators. 67% and 15% of students estimated speed and frequency of wobbling correctly on the first attempt, respectively, with the latter reflecting a more challenging concept. The median speed was 60*μm*/sec (IQR=69.2, N=148), and the wobbling frequency were 0.5rev/sec (IQR=0.5, N=147). Students appreciated the benefit of visual post experiment analytics, with a median feedback score of 9 (highest score *“very useful”*, IQR=3, N=52) and remarks like, *“I also like the fact of being able to download the videos so I could stop them when I was trying to find something or notice anything I did not see at first in the experiment.”*

#### General Design Principles

Image data is information rich. Extracting, augmenting, and visualizing information from the raw data, also in redundant fashion (e.g., different ways of symbolizing the stimulus intensity), enables students to effortlessly explore, and furthermore to focus them on specific details. For example, students were already able to measure length, velocity and rotation frequency of Euglena by inspecting the visualization. Such “back of the envelope” measurements based on semi-processed image data also provide a valuable intermediate between pure qualitative analysis and full data processing (see unit #5) without having to use any specialized software or programming.

### Unit 4: Model exploration and evaluation

Unit #4 featured a modeling environment (Fig. 3E) that prompted students to find the best parameter values that fit the model to the real Euglena path and then explore how to accomplish both positive and negative phototaxis. Students were introduced to the relevant sub-cellular structure of Euglena and the mechanistic explanation of Euglena phototaxis, i.e., the coupling of the eye spot with the flagellum (Fig. 2B), which causes rolling around the long axis and side way turning. These activities then also provided a deeper explanation for the wobbling motion analyzed in the previous unit. Students were also asked to go back to the real experiment after the modeling and to report on similarities and differences.

#### Design Rationale

The target of this unit was to focus students’ attention towards the role of the three simple parameters *surge* (forward velocity), *roll* (rotation about its long axis) and *coupling* (sensitivity to light) that are necessary for phototaxis mechanism; similar to bifocal modeling [3], they had to further compare modeling results with the behavior of real Euglena. In contrast to our previous middle-school study [20], the light sequence was pre-programmed and students could not influence the light sequence during the simulation, which enabled them to concentrate on parameter fitting only. A separate controlled study [5] had revealed that without the joystick, students were significantly more systematic with their parameter exploration. Overall, the modeling interface was intended to provide students with a deeper insight into the core mechanism underlying biological phenomenon, which then makes observation of the real noisy data more vivid.

#### Activities and Feedback

We found that 48% (N=77) students were successful in the fitting tasks, i.e., found a parameter set that led to a closely matching swimming path. When comparing the model to real experiments again, 69% of students noted that the behavior of real Euglena changed depending on the light intensity, but not the model behavior: some real Euglena spun in one place (40%), wobbled differently (19%), seemingly increased in forward or rotational speed (15%), or moved towards the light (8%). 31% of students noted variations in real Euglena behavior both within a single cell and in the population. Finally, 15% of students noted that real Euglena lagged in their reaction to light, whereas the reaction of the model was always instantaneous. These observations show that the students recognized subtle yet significant differences that went beyond what was explicitly discussed in the instructions. We argue that these differences became obvious mainly due to the juxtaposition of model and real Euglena. Students rated the modeling activity to be very useful (8.4 ± 1.2 on the scale of 0-9, N=51) and expressed that view in their comments, *“The simulation was very interesting and improved the learning s it was easier to observe the phenomena [sic].”, “I think to learn better a mix of simulation and real videos is the best option.”*

Similar to unit #2, we analyzed how the student approached modeling more deeply: We identified four strategies that students attempted to explore the parameter space from *N* = 1531 modeling experiments. For this, we computed the number of parameter changes between successive runs which can range from 0-3, thus turning a sequence of experimental runs into a sequence of parameter changes (states). We removed all the 0 states from this sequence, i.e., discounted all the experiment repetitions and computed a single Markov transition probability matrix (where *M*_*ij*_ indicates the probability of going from state *i* to state *j*, where *i, j ∈* {1, 2, 3}) by counting state changes in the sequence. This transition matrix encodes the sequence of the modeling experiments run by a particular student. We then extracted four dominant clusters (silhouette score: 0.58 in 0 – 1 range [35]) from the transition matrices of all the students using hierarchical clustering (Fig. 7). Cluster 1 and 2 (Fig. 7) reveals the most efficient strategy as students in these clusters ran only 14 and 14.5 (median) experiments respectively before posting a near optimal solution. Students in these clusters predominantly switched to changing 1 parameter only followed by multiple parameters at a time. Whereas in cluster 4, which proved to be the most inefficient strategy, students ran 25 (median) experiment for the same task and the key difference with other clusters was that the students, in rare case when changing all 3 parameters, the students predominantly followed that up with 2 parameters changes, which is evident from the transition matrix visualization (Fig. 7). Finally, students in cluster 3 mostly changed 2 parameters at a time and required 17 experiments for the task. We did not notice any significant differences in the number of repeated experiments across clusters. These observations are consistent with the smaller scale middle-school studies [20, 5].

**Figure 7:**
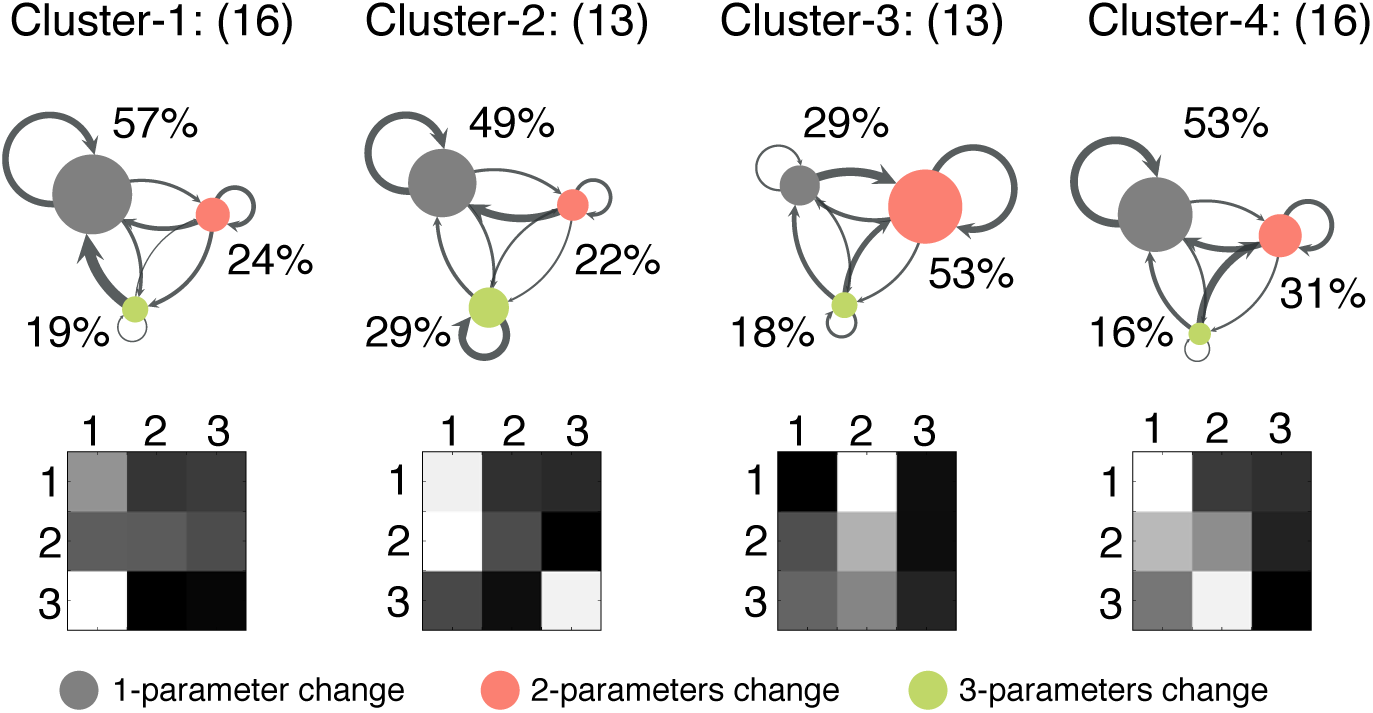
Cluster analysis of parameter fitting strategies for a Euglena model in terms of the transition among the number of parameters changed between successive runs of the model. The transition matrix is shown with a grid where gray scale color coding indicates probabilities of transitions (black=0, white=1). Clusters are arranged from the most efficient to the least efficient strategy (left to right) in terms of how many modeling experiments students ran (median: 14, 14.5, 17 and 25 respectively) before posting a near optimal solution. The numbers in parentheses represent student counts while the percentages are net probabilities of the states.

#### General Design Lessons

The bifocal modeling framework [3] that is centered on the simultaneous comparison of real system and model behaviors provides a productive way for students to both understand the key properties of the real system through the model, yet recognize its subtleties by seeing how real and model behavior diverge. Especially for biological systems that exhibit quite complex behaviors early on, even simple models that capture only the overall tendencies in behaviors can provide powerful lenses to better understand the system under study and appreciate its complexities. In particular mechanistic models, i.e. models that provide mechanistic explanations of the underlying biological phenomena, can form the basis for insightful discussion and investigations.

### Unit 5: Quantitative data processing and analysis

Unit #5 engaged students in the process of exporting the numerical data into a spreadsheet, graphing that data, and interpreting the graphs. Students first worked through a highly scaffolded example to analyze how the Euglena speed depends on Light-On versus Light-Off stimuli, which has at best a weak effect, (Fig. 8, purple trace). Then the students were asked to perform a similar analysis on their own, but now to determine graphically whether the average velocity vectors in x and y changed with directional light stimulus from the LEDs (Fig. 8, pink and cyan traces). Here the students used a data set where the velocity of each cell was already decomposed into its cardinal directions, i.e., Vx and Vy. Depending on the direction of the light origin, one velocity component would average to zero, and the other would be either negative or positive. Students had to design and run a new set of experiments to generate the data for this analysis. Initially, we had intended to teach units #4 and #5 in reverse order, but based on user feedback during pilot studies the interactive modeling activity seemed better before this more complex data analysis.

**Figure 8:**
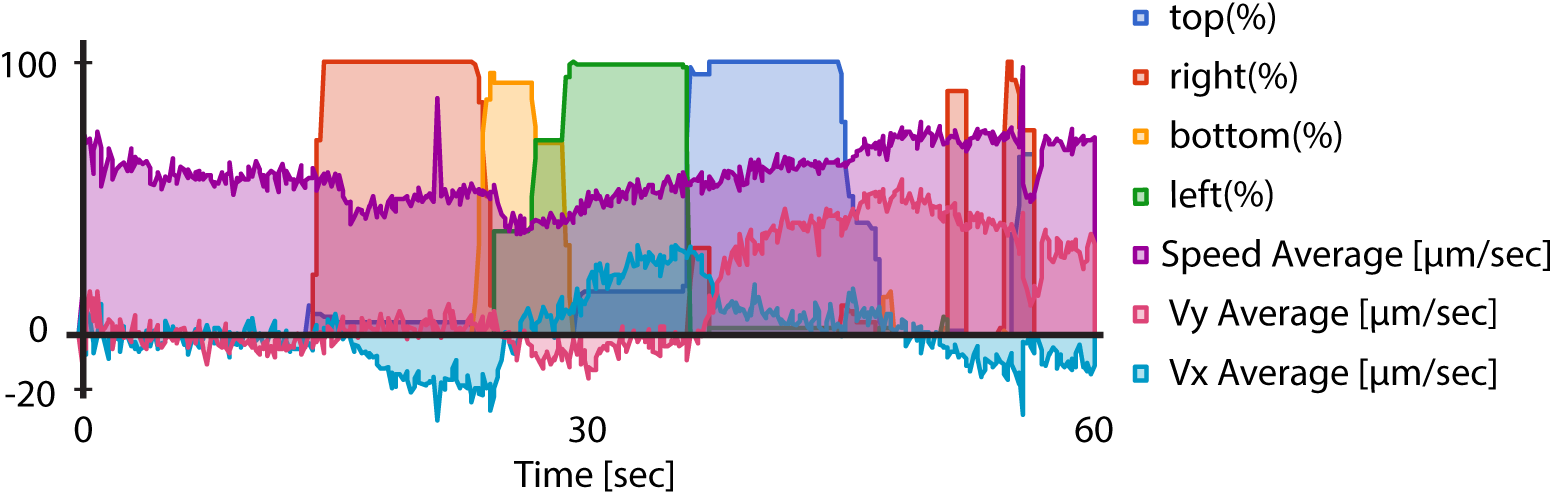
Graphing examples by a student. The light intensity of the four LED is given (top, right, bottom, left). The average Euglena speed (purple) remains stable over time. The average velocity components in x and y (cyan, red) are clearly either positive or negative or zero depending on the light direction.

#### Design Rationale

The purpose of the data analysis component was to export all numerical data in a way that is approachable across a diverse set of audience - from middle school to graduate level students. Through multiple pilot studies we evaluated different data formats, JSON, with corresponding higher level APIs for Matlab and Python. Each had its challenges: Matlab is not free, and the required programming knowledge for Matlab and Python heavily shrinks the potential audience for a large scale online course. Ultimately, we found that the best solution is to export all data to MS Excel *xlsx* format (Fig. 3E), which can also be imported into the Google Spreadsheet, which has built-in advanced statistical and graphical functions. Google Spreadsheet is freely available online, many K-12 teachers are using it with their students, and users in general are familiar with at least some spreadsheet program. However, the row-column spreadsheet format is not as expressive as JSON. In order for the student to be able to generate sophisticated plots with a simple row-column selection and a click of a button, we exported multiple *xlsx* files, each with a suitable data rearrangement and pre-computation, e.g., the population average velocities along the cardinal directions for each time point were already pre-calculated. We also exported the data in the single generic JSON file to enable more advanced students to do more complex analyses.

#### Activities and Feedback

Among the 44 students who voluntarily provided us with the links to their Google Sheets, we analyzed 14 from session 3. We found that 6 students ran the appropriate experiments (i.e., light intensity and duration was such that enough cells exhibited proper responses that could be read off the graph), and also produced the proper graphs including formatting and axis labeling. Another 8 students had proper graphs but their light stimulus protocols had not been optimal. This unit was certainly more complex and challenging than all the previous ones as indicated by its success rate and the overall time the students spent on this unit (on average 1h), and expectedly - due to inherent noise with real biology, some students expressed challenges with the data analytics, *“In general the course explain itself clearly but in some parts like when it’s about the graphics in google sheets and some observations of it, it was a little challenging to understand how I should be able to contrast my own experiments with the examples that are given in the course [sic].”*. However students found this data analytics exercise overall very useful (8.2 ± 1.22 on the scale of 0-9, N=51) with comments like *“I also liked the clear instruction on how to develop useful graphs!”*.

#### General Design Principles

Data formats and analytics UIs should be easily accessible to a broad audience (Google Sheets, CSVs and Excel formats are usually good options). At the same time complex data manipulation should be enabled for more advanced investigations (single generic JSON files and processing in Python seem suitable). We suggest to provide both types of approaches, furthermore to make the activities even more accessible by already pre-processing and pre-formatting the data significantly.

### Unit 6: Open and self-guided investigations

In the final activity unit, students were led to carry out a self-guided research activity, where they proceeded through the main parts of the inquiry cycle while applying all or most of the previously used tools. We prompted students to make an observation (specifically one that had not been stated by the course material previously), and transform this observation into a testable hypothesis with experimental designs. Students were then encouraged (optionally) to pursue the actual experimentation, analysis, and interpretation.

#### Design Rationale

We holistically combined all the UI components (experimentation - both *live* and *batch* - visual analytics, data analytics, and modeling) under the same platform to provide an end-to-end system that allowed students to perform scientific activities remotely over the Internet on real biological samples. In the previous units, we walked students through the key scientific processes and introduced all the UI components to accomplish various scientific tasks. The *batch* experimentation provided students with a key research tool to submit many controlled experiments to be executed offline. This holistic integration of all the UI components, especially with attention given to making the overall system approachable to a diverse audience allowed us to now encourage students to undertake their own research using authentic scientific practices. We made the final project activity optional as an open ended investigation could take significant time, while we wanted to enable students to complete the course within one week.

#### Activities and Feedback

Students made several meaningful observations, some of which were also formulated as a causal clause (*“It seems that when two Euglenas crush, their velocities change.”*), others were more observational (*“Some spin like a gyroscope and others roll. They seem so random.”*). Interestingly, some observations have been published in the recent literature, e.g., *“Many Euglena appear to take at least 2 seconds to move when exposed to sudden intense light.”*, an effect described as transient freezing [30]. This demonstrates that this course and cloud lab enables students to make discoveries equivalent to true science.

65 (20%) students formulated a significant number of distinct and testable hypotheses, 42 (13%) of which were phrased as one variable depending on another (we did not explicitly tell them that this would be a good strategy.): *“The fewer the Euglena in the container, the faster they respond to the light.”*; *“It looks like the rotational speed increases when the light intensity increases.”* We characterized all of the meaningful hypotheses based on *“If the independent variable is (increased, decreased, changed), then the dependent variable will (increase, decrease, change).”* We identified more than 10 classes of both independent and dependant variables each: Independent: Light intensity (on/off, threshold), light direction (two vs. one side), exposure time/minimal time of illumination, cell size, Euglena density, Euglena crashing into each other, different online microscopes; Dependent: Aggregation, stay at one place, directed movement, spinning, spinning frequency, rotational speed (frequency), speed, response time, delay, frequency of cell-cell touching, behavior, behavioral transition, synchronization, activation, interaction between Euglena. Other suggestions did not fall into these categories, such as testing for a correlation between mean and standard deviation of the speed. Hence, well over 100 hypotheses could be generated and tested with the platform, as constrained by the stimulus and observation space, which opens a large possibility space for learners to carry out versatile and self-driven inquiry projects.

21 (6%) students attempted this optional integrated research activity, 15 (5%) of them did it in a meaning-ful way completing all phases of the self-guided investigation; the two following examples serve as illustration (Fig. 9): One student observed that Euglena only reacted at light levels of 50% and higher and decided to investigate what percentage of Euglena move away from the light in response to increasing light levels. The student programmed an experiment in *batch* mode in which light levels systematically increased in steps (Fig. 9A) and reported: *“As the light level increased, movement across the y axis doubled, whereas the x axis stayed consistent.”* The other student observed that Euglena were not always responsive to light and hypothesized that the cells were desensitized or *“exhausted”* by the stimulus, especially after repeated stimulation. The student then designed and ran a batch experiment with a single light-on step (Fig. 9B) and found that the magnitude of velocity away from the light source increased over time but did not desensitize or exhaust the Euglena. Hence the student disproved his hypothesis, but correctly noted that an experimental setup that allows to run experiments for longer than one minute might have helped answer the question better.

**Figure 9:**
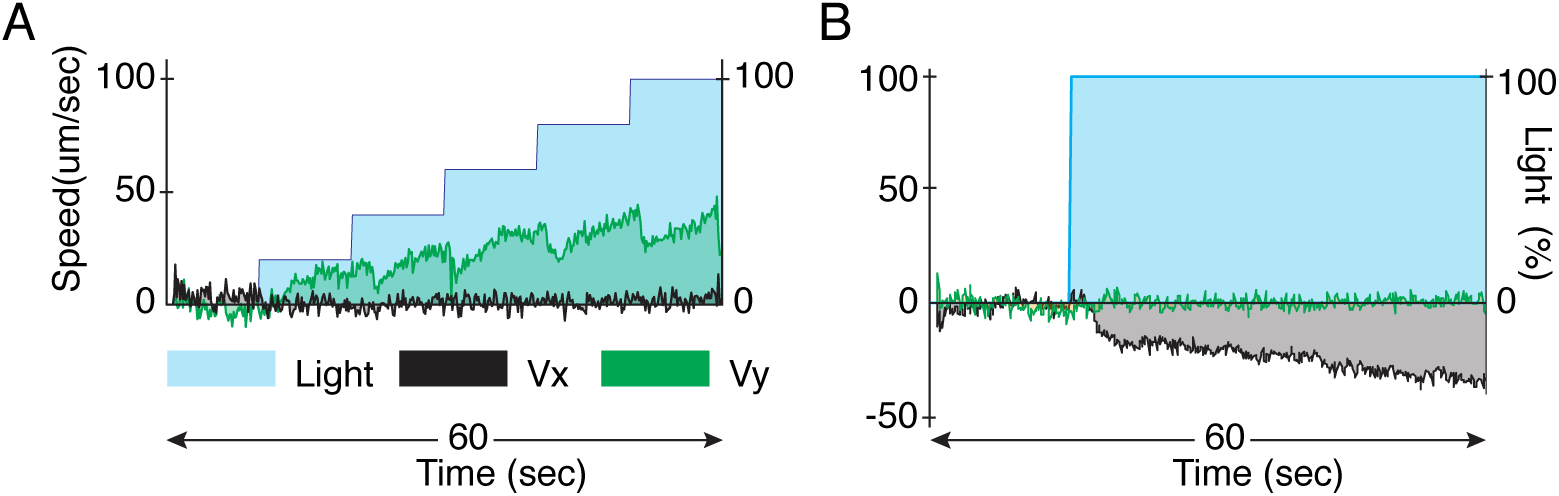
Self-guided student projects. A) Testing the strength of Euglena response in response to increasing light levels (velocity component Vy in green vs. light intensity as blue step trace). B) Testing whether Euglena desensitizes after prolonged light stimulus

#### General Design Principles

To foster inquiry based learning that is in line with NGSS [10] and NRC [39], a cloud lab should integrate all the technology and HCI modules within the same platform in order to reduce switching cost from one inquiry phase to another and reduce as much as possible extraneous technical work that might be a roadblock for the inquiry work (e.g. handling complex data files.) The course should walk students through every phase, introduce the relevant technology in each phase, make sure the students appreciate and understand how to deal with the inherent natural (biological) variabilities in real experimentation (e.g., repeat experiments on multiple instrument and several times) before encouraging students to embark on self-guided research.

### Units 1-6: A case study for the whole course

So far we have described each course unit #(1 - 6) and the relevant design principles independently. We now narrate a case study of a single student’s journey through the course to provide an overall view of how this scaffolded course design, together with the interactive cloud lab, enabled key phases of inquiry-based learning through the Internet. We note that there a variety of ways students approached this course, and a significant portion did not finish it, but we consider the following as a best case scenario given the current course layout, cloud lab, and user interfaces (Fig. 10). This student is a high-school teacher who ran 13 real experiments (9 *live* and 4 *batch*) and 12 modeling experiments in order to complete the course in 3h. She initially ran 5 experiments, including one *batch*, before she noted that Euglena moves faster upon light stimulus but the direction of motion was not clear. Later she executed 3 more experiments on different online microscopes to experience the biological and system variability. By then, she recognized the response to light but the direction of Euglena motion was not as obvious yet to her. She then executed 12 modeling experiments to fit the phototaxis parameters to the correct values, during which time she also executed real experiments to compare with side by side. Such seamless switching between different modes of experiments were possible due the holistic integration of the various components for inquiry-based learning under the same platform. She then ran 3 more experiments before formulating her hypothesis that *“euglena respond when light intensity is above the threshold of* 50%.*”* To test this hypothesis, she ran 2 more carefully designed *batch* experiments in which she ramped up light intensity (top LED only) by 20% every 10s starting from 0% (Fig. 9A). She analyzed the experimental data, which revealed that Euglena swarm velocity along the vertical direction increased with the increasing light intensity while the horizontal component remained constant around 0. This observation confirmed to her that Euglena exhibits phototaxis, which is dependent on light intensity. She voluntarily shared her data plots with us through Google Docs (Fig. 9A), which indicated sound scientific analysis and communication.

**Figure 10:**
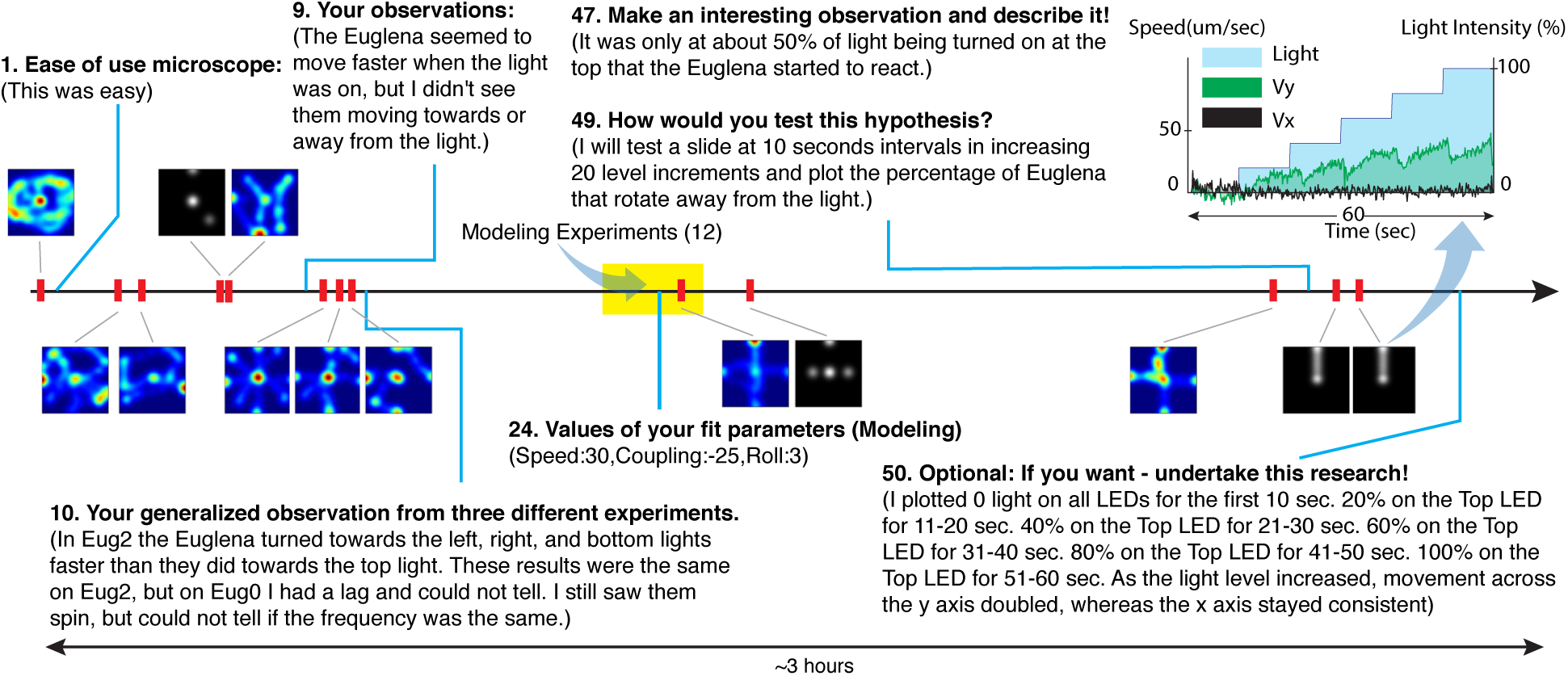
Case study of the activity timeline of one student in the course (same student as in Fig. 9A). Numbers refer the task number in the course (out of 71 tasks total). The colored and grayscale images depict the stimulus pattern of *live* and *batch* experiments, respectively.

### Unit 7: Summary and Reflections

The final unit summarized the course content for the students, provided a set of test questions, and collected overall feedback, which we now discuss with a specific focus regarding some central questions:

#### Accomplishment of course goals

The main goal of this course was to enable students in large numbers to enact the key components of scientific inquiry through the Internet, which was successfully achieved by a significant number of participants (see Fig. 10 for a successful example). The completion rate of 33% is comparably high for MOOCs. We are aware that our course is relatively short compared to a typical MOOC, which likely positively affects the completion rate, but large online courses have considerable dropout rates even within the first few interactions. In comparison to the typical participation profile in open online courses, a 33% completion rate seems promising, especially considering that the course required much more than simply passively watching lectures and taking quizzes. We attribute this high retention rate to the inherent active and live nature of the experimentation in addition to the versatile course content and intuitive interfaces, all of which underwent significant user testing and iterations prior to the course release. Overall student feedback (see below) also speaks to that interpretation. Students self-assessed to have learned about how microorganism interact with their environment *“moderately”* (2.9 ± 0.7, scale of 1-4, N=29), to have learned *“somewhat more than expected”* (3.9 ± 1.0, scale of 1-5, N=29), and students also expressed that they would like to pursue this topic further *“very much”* (5.8 ± 1.1, scale of 1-7, N=24). While this is promising, we plan to perform a thorough analysis of the actual learning outcome in the presence of such a cloud lab as a separate followup study in the future.

#### Changes in attitude towards science

We asked four questions before and after the course to assess students’ attitudes towards science; N=15 students responded pre and post. (These questions were introduced only from session 4 onwards. These 15 students represent about one third of students that completed these sessions.). Answers were on a scale of 1-9 (not at all - totally): *“Science is interesting”* (pre=7.5 ± 1.4 → post = 7.7 ± 1.8); *“I know what it is like to be a scientist”* (7.7 ± 1.3 → 7.4 ± 1.7); *“Ordinary people can be scientists”* (8.7 ± 0.6 → 7.5± 2.0, *p <* 0.05); and *“I can imagine myself as a scientist”* (6.7 1.8 8.5 1.3, *p <* 0.005). The answers to the first two questions were at a high level and did not change significantly. We acknowledge that these responses could in part arise due to a self-selection bias, i.e., students who had a positive attitude towards science were the ones who completed the course and also responded. However, even within this small subset, the changes in the attitude in the third and the fourth questions interestingly revealed that students perceived themselves of being a scientist more than before, though at the same time felt science was more challenging than they initially had thought. Such viewpoints suggest implications that we aim to study separately in the future.

#### Students overall feedback

Students liked the course and rated it as interesting and having the appropriate level of difficulty. All rated their overall experience between *“very”* and *“extremely positive”* (6.3 ± 0.6 on a 1-7 scale, N=34); difficulty was between *“neutral”* to *“somewhat easy”* (4.6 ± 1.1 on a 1-7 scale, N=31); guidance level was leaning towards *“right amount”* (2.8 ± 1.1 on a 1-5 scale, N=34). Students ranked the various lab activities (scale of 1-9, N=52), with *“being able to stimulate cells with light in realtime”* (8.4 ± 1.5), *“modeling”* (8.4 ± 1.2) and *“download your own data and process and graph”* (8.2 ± 1.2) among the most interesting.

#### What students liked

The student feedback about what they liked captures the key features we intended to reach with this online lab course: (1) Value of interactive and remote microscopy (*“The way we could conduct experiments remotely was very cool!”*); (2) Performing deep scientific inquiry (*“Feeling like I was part of real research”*; *“developing a scientific approach to study things”*); (3) Learning biological content (*“how I was able to see diferences [sic] in the behavior of the Euglena”*; *“Learning new things about a microorganism”*); (4) Synergistic integration of different activities and HCI instruments (*“Highly interactive methods of using microscopes, movies, spreadsheets instead of dull passive theory on the characteristics of Euglena.”*); (5) Appropriate course design, content, and length (*“Good amount of material for a short course.”*; *“I liked the emphasis placed on the scientific method.”*; *“I was able to prove myself I was going trough an investigation and how it was going from the easy things to some challenging”*; (6) Lowering access barriers (*“Being from develop country we dont have microscope or all the lab equpiments, this facility has provoked my passion to go for higher studies …[sic]”*; *“I was able to show my child a microscope / microorganisms”*); (7) Playfulness, fun, motivation, personalization, and feeling ownership (*“The course helps make biology fun to learn”*, *“I liked playing around with the online microscope. It was fun looking for phenomenon on your own!”*; *“the way this course has been designed itself is motivating to get through the contents”*, *“The ability to make my own experiments.”*); (8) Advancement beyond what current MOOCs can deliver (*“I like the fact you are using online learning in a different way than most courses.”*, *“It broke this limitation of MOOC courses that they were focused on theoritical [sic] lessons and not the ones requiring laboratory activities”*; *“I have paid some money in Coursera’s lessons and i can say that this was the most interesting lesson i have followed [sic].”*)

#### Suggested improvements

Students pointed out existing limitations and suggested future improvements: (1) extension of experiments (other specimens and organisms; other stimuli beyond white light including chemicals; zoom in and out; better microscope for visualizing the flagellum; longer sessions at the microscope; (2) more explanation and guidance on the data graphing and interpretations; (3) technical improvements such ability to download individual data files instead of a larger compressed file, or the ability to communicate with other students.

#### The value of a real lab vs. a simulation

A central question is whether the effort to provide a real, interactive lab is justified compared to using an interactive computer simulation that is potentially easier to develop and disseminate. The purpose of the present work was not to run a controlled comparative study between a real cloud lab and simulation, instead to provide both under the same platform. We asked the students for their self-reported opinions about the value of real-life experiments over computer simulations, i.e., modeling in unit #4. N=39 students responded and a majority 72% expressed argument in favor of the real lab. 36% explicitly mentioned how simulations may be inadequate at capturing fine details while a real lab provides ground truth data; *“Yes, there should be a real microscope as it is impossible to guarantee that the behaviors of the simulation to be 100% natural/realistic. Using simulations rather than real cells could possibly mean that some unique phenomenon are not discovered.”* The other 36% discussed about the increased course engagement due to the inherent fun and motivating factor that a real lab ushers; *“… using a real microscope is more exciting than using a computer simulation. Being excited is a better motivator for doing the course.”* The remaining students were ambivalent and thought a simulation was adequate for the purpose of this course. A comparative study to measure the actual learning outcome due to simulation versus a cloud lab versus a combination of both (our platform can expose each of settings separately) is an option for future studies.

#### Potential for future integration into K-12 and college education

We extracted feedback from self-identified teachers (K-12 and college, N=12). These teachers generally found the system to be powerful in fostering scientific inquiry due the blending of real biology experimentation with data analysis and modeling; furthermore, filling a current gap regarding the Next Generation Science Standards NGSS [10]. Two teachers explicitly expressed interest in integrating this platform into their high-school biology classes the coming school year: *“I believe that the emphasis on modeling, design, and quantitative analysis would be extremely helpful to AP Biology students, and I would love to try this in my classes.” “I think that this would be a great thing to use with my students … the thinking and feeling of being a scientist would be powerful for them.”* These studies are currently under way. As pointed out earlier, 12 BPUs would be desired to serve students in a regular class concurrently.

## 4 Discussion

### Key design principles for a biology cloud lab

Several UI design rules have been postulated in the HCI literature such as Neilson’s Heuristics [29] and Shneiderman’s Golden Rules [37]. In this paper we discussed six sets of design principles, embedded within each of the first six course units, which we derived through an iterative design process and several pilot studies. These design principles are mostly relevant for a MOOC course with a real backend experimentation lab (even beyond biology), yet some of them could be reformulated in the light of a much broader general purpose Neilson or Shneiderman’s proposition. There are two specific design principles that are unique and particularly important due to the presence of real biology: First, our platform not only mitigated non-deterministic and noisy biological behavior for consistent experimental results, but also provided means to exploit its educational value in the context of the inquiry-based learning. Secondly, biological phenomena are often complex and our design provided a bi-focal modeling platform with a much simplified modeling UI side by side with real experimentation to explain the underlying phototaxis mechanism. These design principles regarding handling of the natural biological variability and bi-focal modeling were key to the success of our biology cloud lab for inquiry-based learning.

### Achievements and implications

We successfully deployed a real biology lab that enables authentic inquiry-based learning for life science in an online learning environment at scale:

(1) We applied the framework of cloud computing to biology experimentation labs, which is beyond simply putting large numbers of microscopes online. We demonstrated that this technology works robustly, can scale linearly to large user numbers (30,000+ experiments/week on 6 BPUs, which represents a two-fold capacity boost from our previous work [20] due to software improvements regarding BPU handling), and at low cost (< 1 ct/experiment) as the BPUs including their biological material require low maintenance effort, and where each BPU is hot-swappable while the system overall remains operational. It is important to realize that this adaptation of a cloud computing-like architecture [16, 38] is the key to enable real science labs at scale (not only in biology) and constitutes a crucial innovation of our work. Our previous work [20] showed that simply “adding microscopes” (or other experimentation devices) to an online course delivery platform does not scale gracefully without a mechanism for automatic instrument and biology health monitoring (Euglena responsiveness to light, cell count, motility), and substantial engineering to perform load balancing in the backend. Issues of consistency of experimental results, maintenance, reliability and redundancy can greatly hinder the learning experience, and our system addresses those concerns successfully by (i) having many BPUs to run experiments on, (ii) automatically monitoring and routing students to the healthiest available BPU (i.e. reducing biological variability) but also giving users an option to self-select a BPU of choice (to expose students to variability on purpose), and (iii) adapting dynamically, e.g. re-routing users, to BPUs and other server failures [20].

(2) We implemented a scalable form of the *interactive biology* paradigm [34, 27, 22] to go beyond what is currently possible and what is the standard for microbiology education: Instead of passively observing through a microscope, students can now interact with cells in real time by applying light stimuli and see direct cellular responses. This also results in rich qualitative and quantitative data (i.e., complex time laps movies as well as automatically tracked swimming paths of many cells), enabling versatile forms of simple all the way to complex data analysis. This contribution is applicable for both presential (face-to-face) and online learning environments.

(3) We integrated a set of HCI modules and activities that realizes the NGSS view of inquiry-based learning at scale, which has not been done with other existing remote labs previousley [18]. This system enabled students to engage in relevant phases of scientific inquiry within a consistent user interface, within which the data was transported and processed in between the different modules to assure students’ focus on the inquiry process and not on extraneous aspects of data handling. The user-friendly yet powerful data-handling formats and software interfaces as well as the short duration of experiments made it possible for students to progress from simple activities all the way to self-driven generation and testing of many possible hypotheses - also given the various possible light stimuli and the information richness of image data. This is in alignment with the *low floor, wide wall, high ceiling* paradigm of constructionist learning tool kits [33]. Students voiced their appreciation for real labs and the course overall, furthermore indicated positive attitude changes towards science.

(4) We converged on key design features (technology and courseware) to not only mitigate the noisy biological behavior (which is inherent to all biological systems), but to actually exploit its educational value. We provided ample opportunities for the students to repeat experiments on different setups and pay attention to variability, furthermore make them recognize the difference between the determinist model and the real biology (biofocal modeling [3]). Recognizing this variability may also increase student interest.

(5) From this work we extract a number of general design principles, discussed in context of the course units in the preceding section, which would extend to other STEM courses with real cloud labs in the future: (i) Students should feel part of a real experimentation environment (e.g. the organisms are real), (ii) have means to initially interact with the underlying phenomena playfully for an intuitive understanding, (iii) are able to execute controlled experiments in batches, (iv) interpret experimental results with minimal effort through visualization, and analyze the data with accessible and familiar tools, (v) understand the mechanism of the underlying phenomena in a noiseless simulated environment and juxtapose the findings with real, noisy experimentation (bifocal modeling [3]), and (vi) are able to test various hypotheses using the system in a self-guided exploratory manner that may go beyond the lesson scope of the course.

(6) We also found that the tight co-development and integration of science activities, biology content, and user interfaces (instruments) is key (Table 1). This also requires many course iterations with small focus groups early on, furthermore the weekly wrap around of new course offerings. Using this approach we were able to deliver and test 5 significant course iterations into 2 months.

### Future work

There are a number of important avenues for future research and development with this cloud lab and course: (1) Refine and test the course content for specific relevant learner groups, such as middle and high-school biology, ultimately paving the way for usage by potentially millions of students annually. (2) Include other relevant scientific practices such as collaborative team work or model building (rather than just parameter exploration) activities. (3) Have participants do more complex projects all the way to geographically-distributed team projects, including sharing experimental data among groups of users, or reanalyzing other students’ data. (4) Explore the potential for citizen science, or even let professional scientists work on the platform. (5) Utilize these platforms for deeper analysis using learning analytics to aid instructors and educational researchers. (6) Extend the platform to other experiment types (other light colors, other organisms, different microbiology experiments). (7) Update the BPU performance protocol, such as automatic LED brightness adjustment for optimal negative phototaxis response and feedback is provided to users on “current instrument quality.”

### Conclusions

In summary, we successfully deployed an open online course with an integrated biology lab in a scalable manner. Students could engage in the core activities of scientific inquiry while interacting with living cells, which goes significantly beyond current educational practices of passive observation through a microscope or using computer simulations or animations; instead the lab automation and ease of data collection and analysis leads to easier logistics and extended lab time for students when working from home. The inherent capabilities for collecting automated learner data and using learning analytics techniques, and the different interaction modalities within the same platform open up interesting research avenues for researchers in education and HCI. This high-dimensional discovery space together with positive user responses regarding their scientific self-efficacy also suggests the opportunity to not just “massify” science labs, but to actually democratize complex scientific practices. This technology could arguably be adapted to K-12 education for millions of users annually in the US and worldwide, filling an unmet need as mandated by the NGSS [10] and other national initiatives.

## 5 Acknowledgements

We are grateful to the members of the Riedel-Kruse and TLTL lab at Stanford University, D. Gilmour, and the involved teachers and students. This project was supported by the NSF Cyberlearning grant (#1324753), NSF awards IIS-1216389, OCI-0753324 and DUE-0938075, furthermore graduate fellowships to Z.H. (SIGF) and E.B. (SGF).

